# Heritable plant phenotypes track light and herbivory levels at fine spatial scales

**DOI:** 10.1101/210765

**Authors:** P.T. Humphrey, A.D. Gloss, J. Frazier, A. C. Nelson–Dittrich, S. Faries, N. K. Whiteman

**Author notes:** Equal contribution (P.T. Humphrey, A.D. Gloss). Author for correspondence: N.K.W.

## Abstract

The biotic and the abiotic environment play a major role in shaping plant phenotypes and their geographic distributions. However, little is known about the extent to which plant phenotypes match local patterns of herbivory across fine-grained habitat mosaics, despite the strong effect of herbivory on plant fitness. Through a reciprocal transplant-common garden experiment with clonally propagated rhizomes, we tested for local phenotypic differentiation in bittercress (Brassicaceae: *Cardamine cordifolia*) plants collected across an ecotonal habitat mosaic. We found that bittercress in sunny meadows (high herbivory) and shaded understories (low herbivory) have diverged in heritable growth and herbivore resistance phenotypes. The expression of these differences was habitat dependent, mirroring patterns of adaptive divergence in phenotypic plasticity between plant populations in meadow and understory habitats at broader geographic scales, and showed no evidence for a constraint imposed by growth–defense tradeoffs. Most notably, plants derived from shade habitats exhibited a weaker shade-induced elongation response (i.e., shade avoidance syndrome, SAS) and reduced resistance to herbivory, relative to plants derived from sun habitats, when both were grown in shade common gardens. Greenhouse experiments revealed that divergent SAS phenotypes in shade conditions were expressed in offspring grown from seed as well. Finally, we observed partially non-overlapping flowering phenology between habitat-types in the field, which may be at least one factor that helps to reinforce habitat-specific phenotypic divergence. Altogether, our study illuminates how a native plant may cope with overlapping biotic and abiotic stressors across a fine-grained habitat mosaic.

## INTRODUCTION

In species distributed across environmental gradients, individuals often exhibit phenotypes that track the local environment (Linhart and Grant 1996; Galloway 2005; Richardson et al. 2014). However, our knowledge regarding the conditions under which these patterns arise and persist has been disproportionately shaped by studies at course-grained rather than fine-grained spatial scales (Richardson et al. 2014). This bias has likely been shaped by the expectation that gene flow among interspersed habitat patches is generally a strong homogenizing force, preventing the establishment of habitat-associated phenotypic and genotypic variation at fine-grained spatial scales (Haldane 1930; Lenormand 2002). Although there is growing evidence that heritable phenotypes track habitat mosaics at fine-grained spatial scales [i.e. *microgeographic* phenotypic divergence; (Richardson et al. 2014)] we have a limited understanding of the molecular, ecological, and evolutionary processes that facilitate and maintain this variation in nature.

Defoliation by insect herbivores exerts strong selection on plant phenotypes (Louda 1984; Prasad et al. 2012; Agrawal et al. 2012). In mustards (Brassicaceae), polymorphisms in genes that modify defensive chemicals underlie adaptation to local herbivore communities (Prasad et al. 2012; Zust et al. 2012), and the magnitude of geographic divergence at such loci is extreme compared to loci across the rest of the genome (Brachi et al. 2015). However, much of the strongest evidence for local adaptation to herbivory is from populations separated by large distances, on the scale of hundreds to thousands of kilometers (km). Although plants were the focus of pioneering research on variation in species distributions (Greig-Smith 1952) and phenotypic and genetic differentiation [reviewed in Linhart (1996)] across fine-grained habitat mosaics, relatively little is known about the relationship between anti-herbivore resistance traits and environmental herbivory pressure at microgeographic scales. A few studies have demonstrated that heritable divergence in herbivore resistance or defensive traits can correlate with variation in herbivore pressure at spatial scales ranging from a few km to only 500 m (Galen et al. 1991; Sork et al. 1993; Pellissier et al. 2014; Dostálek et al. 2016; Sato and Kudoh 2017).

The ability to mount strong defensive phenotypes in habitats where herbivores are abundant, either through phenotypic plasticity or through habitat-specific divergence in loci mediating resistance to herbivory, might be impeded by negative phenotypic correlations between defensive traits and those involved in response to other stressors. While the presence of multiple stressors can select for the coordinated expression of multiple inducible plant traits (Boege 2010), the response to selection may be limited by negative phenotypic correlations and insufficient genetic variance (Agrawal et al. 2010). For example, phenotypic investments in herbivore defense reduced stress tolerance in the mustard *Boechera stricta*, and this negative correlation itself may promote the patchy distribution of plants into habitats that do not impose a fitness cost arising from negative trait correlations (Siemens et al. 2009; Siemens and Haugen 2013; Alsdurf et al. 2013). Negative correlations between growth and defensive traits may impose a particularly strong constraint, and the question of how plants “solve” the dilemma of altering growth to compete for light *and* resist enemies in the face of growth-defense trade-offs has been the subject of intense study (Herms and Mattson 1992; Cipollini 2004; Züst and Agrawal 2017).

Herbaceous plant species abundant in both open habitats and deeply shaded forest understories, which often face distinct levels of both herbivory (Louda et al. 1987) and light competition (Dudley and Schmitt 1995), present an excellent opportunity for testing if both growth and anti-herbivore resistance phenotypes can diverge across microgeographic habitat mosaics, despite the potential tradeoffs between growth and defense. This is because decades of study have yielded clear expectations of which growth phenotypes are beneficial in high and low light habitats. In open and sunny habitats, plants express altered growth patterns in response to shade from herbaceous competitors. This change in plant growth traits and architecture, termed the shade avoidance syndrome (SAS), manifests as elongation of stems, petioles and hypocotyls, apical dominance, and early flowering (Keuskamp et al. 2010) induced by perception of light with a high far red (λ=700– 800 nm):red (λ=600–700 nm) ratio or low blue light levels (λ=400–500 nm). Variation in SAS within species is an important example of locally adaptive plasticity (Van Kleunen and Fischer 2005; Valladares et al. 2007). Herbaceous plant species adapted to forest understories—where shading by forest canopies cannot be overcome—benefit from reduced SAS expression, while conspecifics adapted to open habitats benefit from their SAS expression in response to neighbor shade because neighbor shade can more easily be overcome (Dudley and Schmitt 1995; Schmitt et al. 1995; Dudley and Schmitt 1996; Donohue et al. 2000; Donohue et al. 2001; Bell and Galloway 2008). When light competition and herbivory are both encountered (e.g. in meadows), expression of SAS due to neighbor shade may attenuate herbivore resistance owing to an underlying trade-off between growth and defense (Uriarte et al. 2002; Cipollini 2004; Valladares et al. 2007). Whether herbivore resistance and SAS co-diverge at microgeographic scales, despite the commonly observed tradeoff between the two traits, remains largely unexplored for native plants.

Here, we quantified phenotypic divergence in growth and resistance to herbivory across a microgeographic habitat mosaic in bittercress (*Cardamine cordifolia* Gray; Brassicaceae), a forb native to montane regions of western North America. Bittercress grows in clumps across ecotonal patches with sharply contrasting selective regimes: high herbivory with high light competition from neighboring forbs in open meadow (sun) habitats, vs. low herbivory and shading from the evergreen tree canopy in nearby deeply shaded (shade) habitats with fewer herbaceous neighbors (Louda 1984; Collinge and Louda 1989; Louda and Rodman 1996; Alexandre et al. 2017). One of the primary herbivores of bittercress is a specialist leaf miner, *Scaptomyza nigrita* (Drosophilidae) (Collinge and Louda 1989; Gloss et al. 2014; Humphrey et al. 2014; Humphrey et al. 2016; Alexandre et al. 2017). When given a choice in laboratory experiments, *S. nigrita* prefer to attack shade-derived compared to sun-derived bittercress, but abiotic factors over-ride this preference: *S. nigrita* adults strongly prefer brighter and warmer sunny habitats over darker and colder shade habitats, which largely restricts herbivory by *S. nigrita* to sunny habitats (Alexandre et al. 2017). For bittercress growing in the shade, one consequence of prolonged exposure to enemy-free space and decreased neighbor shade might be habitat-associated divergence in the investment in, and/or expression of, both SAS and inducible herbivore defenses. Thus, bittercress presents an opportunity to test for microgeographic divergence in growth and defensive phenotypes across habitats, in a context where such fine scale divergence might be constrained by growth–defense tradeoffs and homogenized by inter-habitat dispersal and gene flow.

Through reciprocal transplant–common garden experiments in the field and greenhouse, we tested if bittercress from deep shade habitats differed in growth phenotypes (including those reflecting SAS) compared to their sun-derived conspecifics. This design allowed us to measure the effect of phenotypic plasticity caused by growth environment (sunny vs. shaded), and whether source habitat impacts the average expression and/or the plasticity of expression of plant growth traits. We coupled these plant growth measurements with herbivore bioassays in which we measured bittercress resistance to *S. nigrita* in the common gardens. Finally, we determined whether flowering phenology differs between habitats in a manner that could facilitate phenotypic divergence by reducing gene flow among habitats.

Altogether, our study illuminates how a native plant may cope with overlapping biotic and abiotic stressors across distinct, fine-grained habitat types. We found that bittercress in sunny and shaded habitats have diverged in heritable growth and herbivore resistance traits. The expression of these differences was habitat dependent and in some cases counter to expected growth–defense tradeoffs. Finally, partially non-overlapping flowering phenology observed between habitat types may be at least one factor that helps to reinforce habitat-specific divergence.

## MATERIALS AND METHODS

### Common gardens in the field

This study was conducted near the Rocky Mountain Biological Laboratory (RMBL) in Gothic, Colorado, USA, between 2011–2014. In 2011 at the RMBL, we conducted a reciprocal transplant-common garden experiment to test if habitat of origin (sun vs. shade) impacted plant phenotypic responses to shading and realized resistance to herbivory. We chose nine sun and nine shade source sites from which to sample bittercress rhizomes for planting in common gardens that were either in meadows (sun) or under evergreen forest canopies (shade) (Fig. S1). At source and common garden sites, we recorded photosynthetically active radiation (PAR) using a light meter (Spectrum Technologies, Inc.), percent canopy cover using a densiometer, diameter at breast height (*dbh*) of the four largest trees within four meters, and latitude, longitude, and elevation (Garmin GPS) (Fig. S2, Tables S1). Source sites significantly differed in PAR, canopy cover, and *dbh* (ANOVAs, all P<0.05) but did not differ in elevation (P>0.05; Table S2).

From each source site, we collected rhizome tissue from 30 ramets and stored them at 4–10°C in low ambient light until planting. Using a randomized complete block design, 540 rhizomes—five from each of the 18 source sites (90 in total per garden)—were planted among six common gardens and spaced 16 cm apart to avoid shading from neighboring bittercress. Gardens were watered every 24–48 h with nearby water from a snow melt stream. Plants were shrouded using fine mesh cloth to prevent herbivory throughout the experiment.

After five weeks, we counted the number of leaves >10 mm and harvested and photographed the largest leaf. We measured petiole length and leaf area using ImageJ (Abràmoff et al. 2004) and leaf mass using a Sartorius CPA225D balance (Sartorius AG, Goettingen, Germany) following oven drying at 65°C for 2 d. Subsequently, we transplanted a single *S. nigrita* larva into the largest remaining leaf. Larvae were collected from mined leaves at Copper Creek, near the RMBL. After 72 h, we removed leaves and measured larval mass as a measure of larval performance (Whiteman et al. 2011; Whiteman et al. 2012), which is an index of realized resistance of the plants. Early instar larvae were excluded to approximately standardize larval developmental stage, but larvae were not weighed in order to minimize their manipulation. We therefore caution that the amount of variance within treatments is likely inflated by variation in initial larval mass. We compared larval mass using two-tailed two-sample *t*-tests of the larval mass distributions between sun and shade source plants in each of the garden types.

### Greenhouse experiment

We conducted a simulated shading experiment under greenhouse conditions using bittercress grown from seeds collected from sun and shade habitats. In August 2012, we collected seeds from bittercress growing in sun and shade habitats near the source sites for the common gardens by tying bags to developing racemes (see Table S3 for collection sites). From the source sites we recorded % canopy cover using a densiometer and latitude, longitude, and elevation of each collection site as above. Seeds were transported to the University of Arizona and surface-sterilized with a solution of 50% bleach and 0.5% Triton-X-100 and stratified on moist filter paper in petri dishes for five weeks at 4°C. Seeds were germinated in the greenhouse in moist filter paper in petri dishes; seedlings were planted in soil in plastic pots as above and randomized to light filter environments to simulate sun or shade habitats. Evergreen forest canopy shade was simulated by growing 14 shade-and 23 sun-source plants under a blue plastic filter (Lee filter 115) that reduces the amount of red light available to plants (elevating the amount of FR:R light) (Runkle and Heins 2001). To simulate sun habitats, 19 shade-and 28 sun-source plants were grown under a control clear filter that does not alter the quality of light (Lee filter 130). When plants reached the 4-leaf stage and the 8-leaf stage, we measured vegetative traits as in the field common gardens. We did not have access to *S. nigrita* herbivores for the greenhouse experiment at to measure realized resistance in this trial.

We conducted all statistical analyses using R v.3.3 (R Core Team 2013). Source site characteristics were compared using ANOVA or linear mixed models (LMMs) using the *lme4* package (Bates et al. 2014). Hypothesis testing on fixed effects was conducted with conditional *F*-tests using denominator degrees of freedom (Kenward and Roger 1997) as implemented in R package *pbkrtest* (Halekoh and Højsgaard 2014). For the field common garden experiment, we modeled garden type, source habitat and interactions as fixed factors and garden number and source site number as random factors. To assess the role of garden and source types on plant phenotypes, we conducted principal component analysis (PCA) on total number of leaves, leaf area, petiole length, and specific leaf area (SLA, *cm^2^/g*). We tested the relationship between garden and source types on PC1 + PC2 using linear discriminant function analysis (DFA) with R package *MASS* (Venables & Ripley 2002). To examine differences in petiole length among sun and shade-derived plants grown from seed in the greenhouse, we used R package *pbkrtest* as above to examine LMMs modeling petiole length at the 4-and 8-leaf stages as a function of environment type, source type, and their interaction as a fixed-effects with source site as a random effect. We separately analyzed petiole length at the 8-leaf stage in the FR:R light greenhouse treatment with source type as a fixed effect using the same approach as above. All field common garden data, as well as statistical analysis R scripts, are available on the Dryad digital repository (doi pending).

### Field surveys of flowering phenology

In 2015, we quantified floral abundance through time in sun and shade habitats in a sample of 240 bittercress plants spread across two sites (401 Trail and Copper Creek). At each site, we designated roughly equal numbers of plants in open sun, intermediate shade, and deep-shade and tracked plant-level flower abundance every 2–3 days for four weeks (between July 8 and August 5 of 2015). From these data, we designated a first, peak, and last flowering Julian day for each stem. In many cases, focal stems flowered before the observation window, and these stems were assigned a Julian date of 1 prior to the first observation day. Plants with no flowers or fruits by the end of the observation window were assigned a no-flowering status and were excluded from the phenology analysis. We also accounted for the presence of developing fruits on each stem, which allow us to designate plants with no *observed* flowers as having already completed flowering by the first observation date, thus distinguishing this class from those plants that had neither flowered nor fruited during the observation window. Many plants had just begun to flower by the end of the observation widow; the ‘peak’ and ‘last’ flowering events for these stems was assigned a Julian date of the last observation day +1. We compared the accumulation of each event across habitats and sites using non-parametric two-sample Kolmogorov–Smirnov tests, which calculates the difference between two empirical cumulative distributions; KS test *p*-values derived from the Kolmogorov distribution (by default) were compared to those obtained by permutation of the identity of the source habitat across data points (1000 replicates).

## RESULTS

### Effects source habitat and light environment on plant growth

In the field, we measured plant growth traits in common gardens to investigate the strength of shade-induced growth phenotypes in bittercress genotypes derived from sun and shade habitats. After six weeks of growth in these common gardens, plants in shade gardens were strongly differentiated from those grown in sun gardens in a PCA using total number of leaves, leaf area, SLA, and petiole length (Fig. 1A). Using PC1 and PC2, which together comprised 80% of the variance, linear DFA correctly assigned 99.2% of plants to garden type, while 61.9% could be correctly assigned to source habitat type. Plants from both source habitats had similarly higher specific leaf area in shade gardens (*F*_1,4.24_ = 46.7, *P* = 0.002; Table 1, Fig. 1B), and plant from both source habitats regrew more leaves by 6 weeks in sun gardens than in shade gardens (*F*_1,4_ = 18.16, *P* = 0.013; Table 1, Fig. 1C.). Means and standard errors of all growth traits for each source habitat in each garden are compiled in Table S4.

**Table 1.** Model results for plant growth traits in sun and shade common gardens.

| Predictor variables | Model statistic | Response variables |  |  |  |
| --- | --- | --- | --- | --- | --- |
|  |  | Num. Leaves * | Leaf Area (cm<sup>2</sup>)†, 1 | Specific Leaf Area (cm<sup>2</sup>/g)<sup>1</sup> | Petiole Length (mm)*, 1 |
| Garden Type | *MS<sup>a</sup>* | 6.68 | 8.4 | 3.40E+05 | 5.923 |
|  | *denDF*†† | 4 | 4.7 | 4.24 | 4.22 |
|  | *F* | 18.16 | 0.92 | 46.67 | 4.56 |
|  | *P<sup>b</sup>* | 0.013 | 0.383 | 0.002 | 0.096 |
| Source Type | *MS<sup>a</sup>* | 5.977 | 0.31 | 3098 | 11.034 |
|  | *denDF* | 15.68 | 17.35 | 16.84 | 18.45853 |
|  | *F* | 15.81 | 0 | 0.42 | 10.54 |
|  | *P* | 0.001 | 0.853 | 0.527 | 0.004 |
| Garden Type X Source Type | *MS* | 0.665 | 81.08 | 16921 | 13.236 |
|  | *denDF* | 324.37 | 264.34 | 260.99 | 265.6573 |
|  | *F* | 1.77 | 9.49 | 2.29 | 13.47 |
|  | *P* | 0.182 | 0.002 | 0.131 | 0.0003 |
| Covariate | *MS* | - | 392.99 | 83500 | 42.23 |
|  | *denDF* | - | 239.9 | 243.08 | 182.03 |
|  | *F* | - | 38.57 | 4.51 | 32.26 |
|  | *P* | - | 0 | 0.0356 | 1.00E-07 |
| Genet X Garden Type | *Var* | 0.0321 | 0.0167 | 17781 | 0.18922 |
|  | *X<sup>2</sup>* | 5.207 | 0.735 | 5.3697 | 3.194 |
|  | *df* | 3 | 3 | 3 | 3 |
|  | *P* | 0.157 | 0.865 | 0.147 | 0.363 |

\*- model results for square root transformed data
†- model results for *ln*-transformed data
†† - denominator degrees of freedom
a - MS from reduced model after testing for fixed and random interactions but retaining fixed interaction term if P<0.1
b - *F* and *P* values from Kendall-Rogers correction for denominator d.f. (see methods)
1 - modeled using leaf number as covariate
2 - modeled using SLA as covariate

**Fig. 1.**
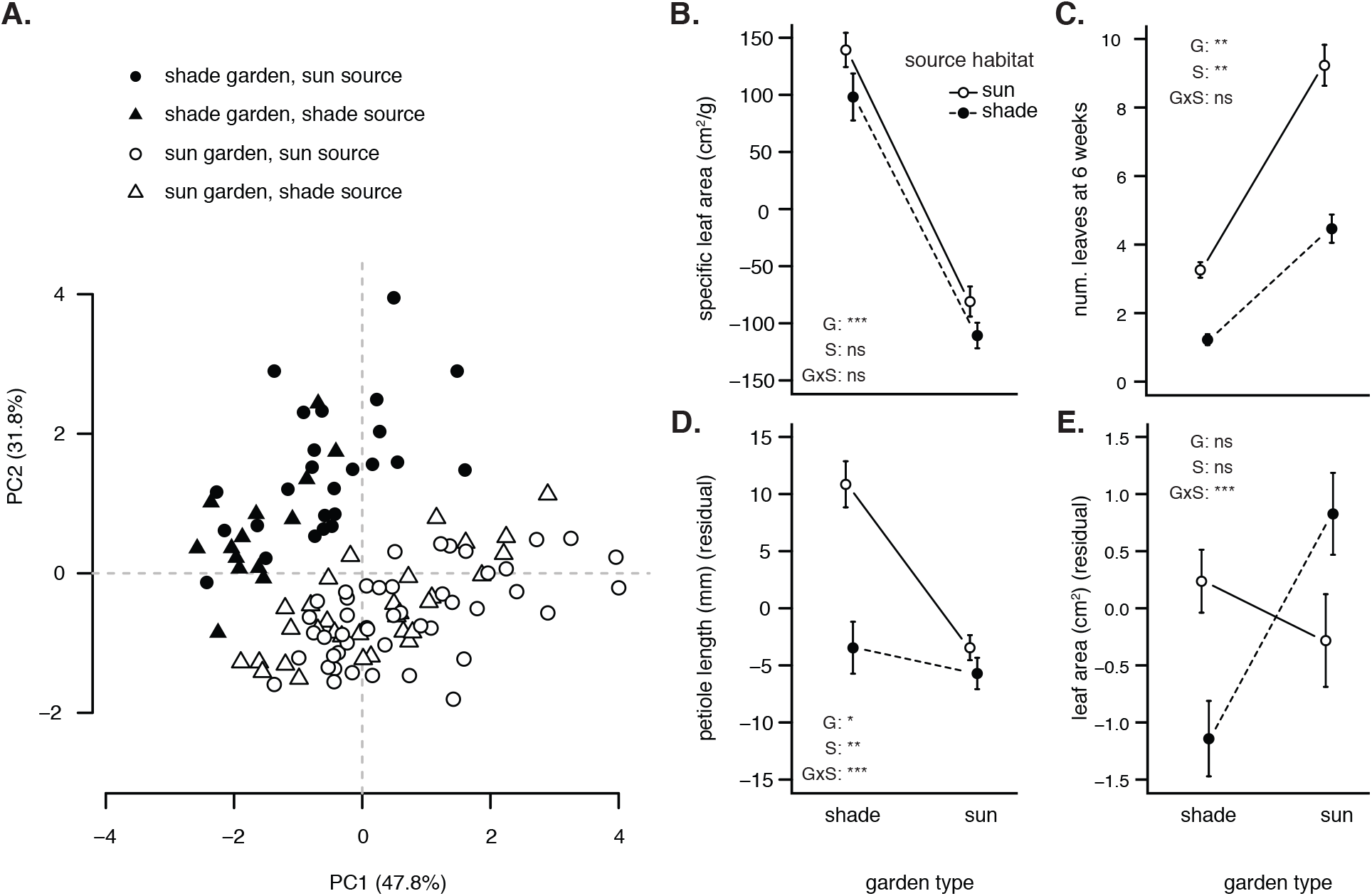
Light environment and source habitat distinguish growth phenotypes of bittercress grown from rhizomes in common gardens. (**A**) Principle components analysis (PCA) of five growth phenotypes (see Methods) reveals distinct phenotypic clusters based on garden type. (**B–E**) Plant traits exhibited a garden type main effect (**B**, **C**), source habitat main effect (**C**, **D**), and interactions between garden type and source habitat (**D**, **E**). Error bars indicate ± standard error. ‘ns’ = *P*>0.05, *0.05>*P*>0.01, **0.01>*P*>0.001, \*\*\**P*<0.001.

In addition to these strong garden type effects, source habitat type affected various individual plant growth phenotypes. Plants derived from shade habitats were smaller than sun source plants in both garden types (*F*_1,15.68_ = 15.81, *P* = 0.001, Table 1, Fig. 1B). Sun source plants had longer petiole length relative to their plant size in both garden types (*F*_1,18.46_ = 10.54, *P* = 0.0044). But the increase in size-specific petiole elongation for sun-derived plants from growth in the shade was far stronger than that for shade-derived plants (*F*_1,265.65_ = 13.47, *P* < 0.001; Table 1, Fig. 1D). This result was recapitulated in the greenhouse, where sun bittercress grown from seed exhibited significantly longer petioles at the eight-leaf stage compared to shade source plants when both were grown under simulated neighbor shade (high FR:R light; *z* = 3.35, *P* = 0.0008, Fig. S3). Leaf area exhibited no main source or garden type effects but showed a strong interaction effect, with the changes in size-specific leaf area being comparable in magnitude between garden types for plants from both source habitats (*F*_1,264.34_ = 9.5, *P* = 0.002; Table 1, Fig. 1E).

### Effects of source habitat and light environment on realized herbivore resistance

In the field, we assayed realized plant resistance to *S. nigrita* larvae by measuring larval mass as a proxy for herbivore performance in bittercress grown in the common gardens. Despite high levels of variation in larval mass across both garden types, shade-derived plants harbored larvae that had a higher average mass after feeding for 24 hours than larvae randomized to any other condition. Specifically, in shade gardens, larval mass was 21.3% higher when *S. nigrita* leaf-miners were reared on plants from shade-derived compared to sun-derived plants (1.59 mg vs.1.31 mg, two-tailed *t*-test, *P* = 0.02, Fig. 2). Within sun gardens, larval mass did not differ between sun-or shade-derived plants (*t*-test, *P* = 0.88, Fig. 2).

**Fig. 2.**
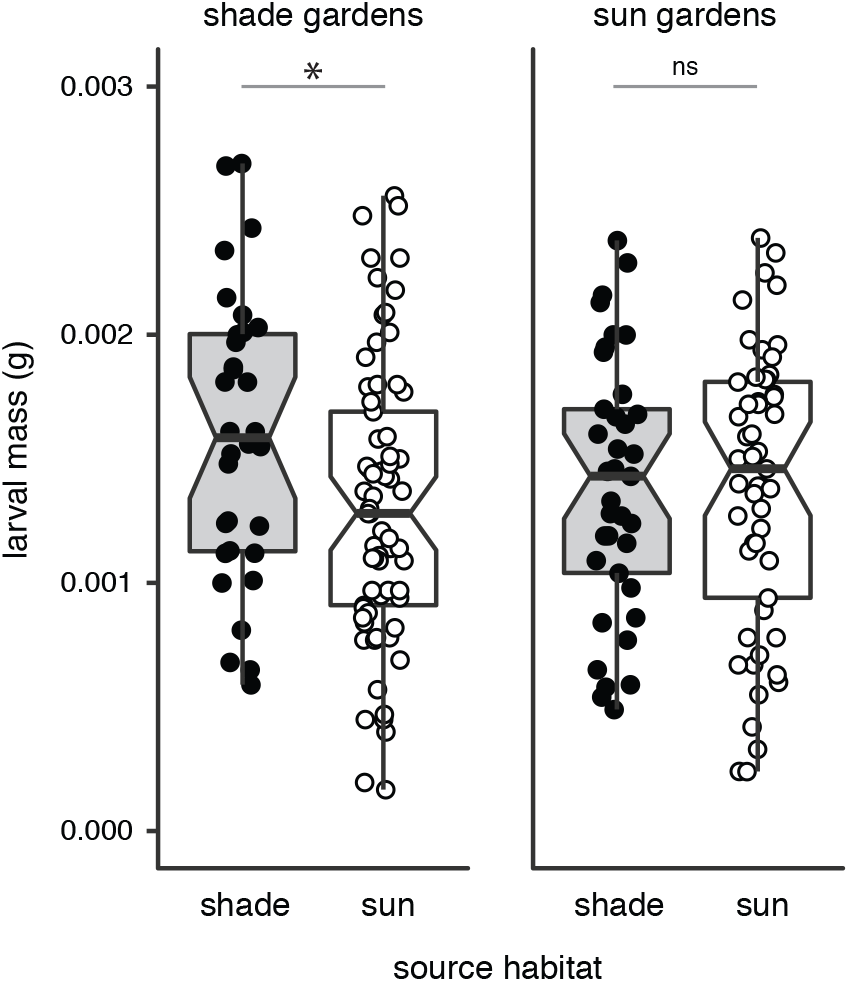
Realized plant resistance to *Scaptomyza nigrita* larvae differed by source habitat in shade but not sun common gardens. *S. nigrita* larval mass was higher on average after feeding for 72 h on shade-derived plants than on sun-derived plants in shade common gardens, while larval mass was indistinguishable between plants from both sources in sun exposed common gardens. Larvae were initially of the same developmental instar but were not pre-weighted (see *Methods*). ‘ns’ = *P*>0.05, \**P*<0.05.

### Effects of source habitat on flowering phenology in the field

Bittercress phenological progression and floral density curves differed both by sampling site and by habitat type. At the 401 Trail site, plants in all habitats exhibited very similar phenological progressions (K–S tests, all comparisons *P*>0.05; Fig. 3A). In contrast, at the Copper Creek site, timing of first, peak, and last flowering was significantly delayed for bittercress growing in intermediate shade and deep shade compared to open sun habitats (sun vs. intermediate or deep shade all *P*<0.05; Fig. 3A). The total density of flowering plants through time strongly overlapped among habitats at the 401 Trail site, while shade habitats exhibited their peak flower density after both the open sun and intermediate shade plants at Copper Creek (Fig. 3B).

**Fig. 3.**
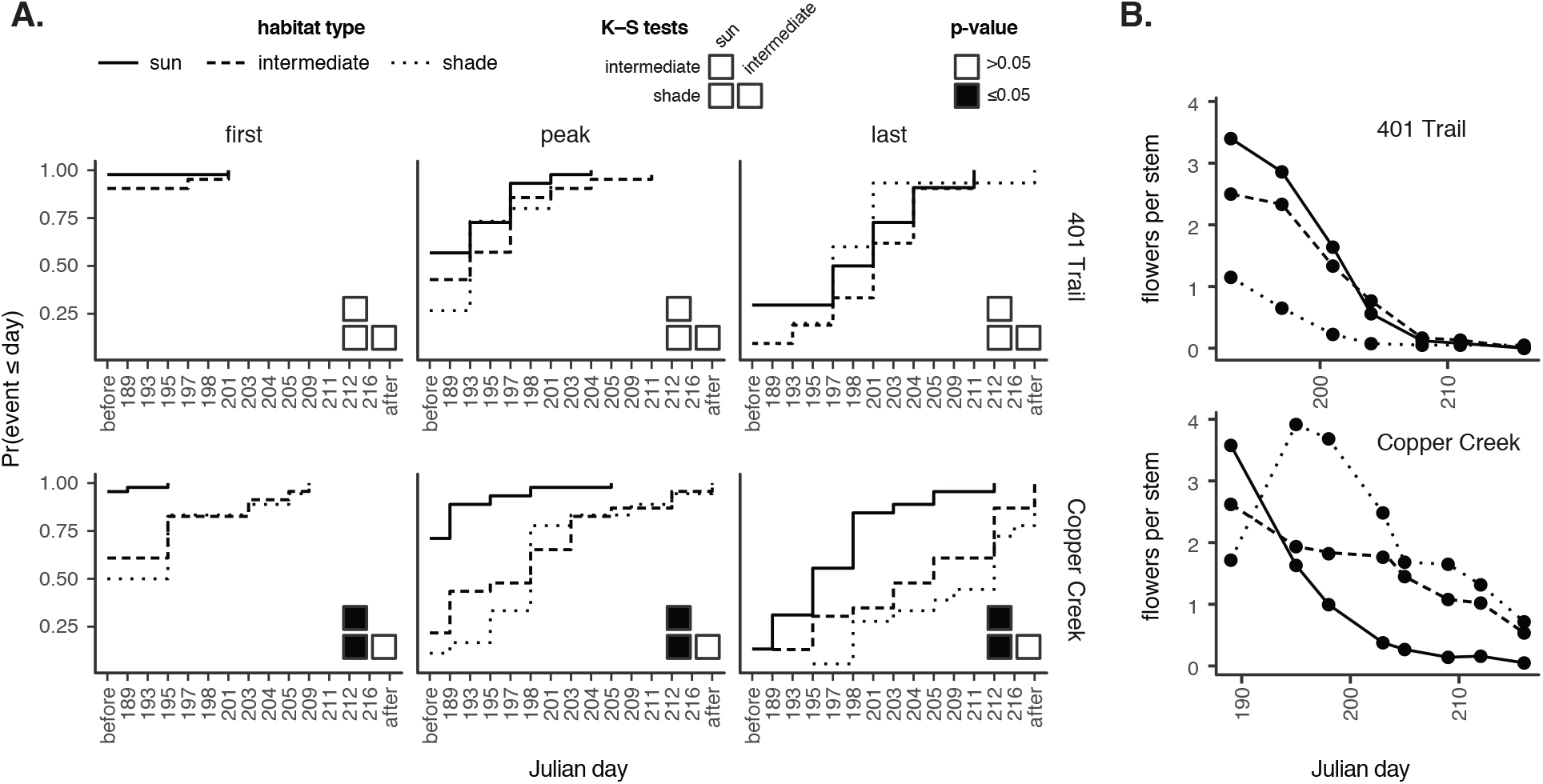
Phenological progression and flowering curves indicate site-specific differences in phenology between open sun and shaded habitats. **(A.)** Phenological progression through first, peak, and last flowering depicted as empirical cumulative distributions indicate faster flowering for sun plants at Copper Creek but not 401 Trail site, as indicated by pairwise two-sample and two-tailed Kolmogorov–Smirnov (K–S) tests. (**B.**) Pooled average flower density through time shows delayed flowering peak in shade habitats at Copper Creek but not 401 Trail site.

## DISCUSSION

### Heritable divergence in plant growth and resistance traits across a fine-grained habitat mosaic

Two notable ecological features reliably distinguish sun and shade habitats for bittercress (Fig. S4), and thus could promote habitat-specific phenotypic divergence. First, herbivory by *S. nigrita*, a specialist leaf-miner that can heavily defoliate stands of bittercress, is primarily restricted to sun habitats (Collinge and Louda 1989; Louda and Rodman 1996; Alexandre et al. 2017). The enemy free space associated with shade habitats would be expected to favor lower resistance to herbivory in plants growing in shade (Agrawal et al. 2012). Second, the growth form of neighboring plants that filter incoming light, either forbs or canopy trees, differs among habitat types. Conspecific and heterospecific forbs grow at high densities in sun habitats of bittercress, and SAS is frequently observed to be adaptive in this context (Van Kleunen and Fischer 2005). In contrast, large evergreen trees filter the majority of light in shade habitats (Table S1). Canonical SAS phenotypes are typically an ineffective means to compete for light in forest understories, and their expression has been shown to reduce organismal fitness (Dudley and Schmitt 1995; Schmitt et al. 1995; Dudley and Schmitt 1996; Donohue et al. 2000; Donohue et al. 2001; Bell and Galloway 2008). We thus hypothesized that sun and shade plants should diverge in their expression of competitive growth (SAS) and resistance phenotypes.

Our reciprocal transplant/common garden assays revealed that heritable plant growth and resistance phenotypes were distinguishable between plants from the shade and the sun habitats. The directions of these differences were concordant with the hypothesis that light competition and herbivore prevalence drive phenotypic divergence among sun and shade habitats. Notably, some of the traits we measured were shaped by an interaction between light environment and parental habitat. In other words, although these phenotypes were plastic, differences in their expression across light environments (i.e., the degree of plasticity) were heritable.

#### Plant growth traits

While light environment of the common gardens strongly influenced plant growth phenotypes, those phenotypes characteristic of adaptive plasticity to shade (i.e., SAS), most notably petiole elongation, differed markedly between plants from sun and shade habitat types (Fig. 1). Rhizomes from sun source sites exhibited strong SAS when grown in shade gardens in the field (Fig. 1D), marked by increased petiole elongation relative to rhizomes from shade source sites grown under the same conditions. We observed this effect both in plants grown from seed derived from the same habitats in the greenhouse (Fig. S3), indicating that this phenotype is inherited by offspring through both clonal and sexual reproduction.

#### Resistance traits

Herbivore resistance is a measure of how well herbivores can exploit a given host resource. We found that *S. nigrita* larvae feeding on shade-source plants gained more mass than larvae feeding on sun-source plants, but this pattern only emerged in shade common gardens (Fig. 2). This finding is consistent with the hypothesis that shade-derived plants invest less in defense when in their natal habitats, where investment in anti-herbivore defenses is likely costly in the absence of herbivory (Agrawal et al. 2012). More sensitive bioassays, as well as additional biomarkers of basal and inducible herbivore resistance, are required to identify the defense phenotypes responsible for the observed difference.

### No evidence that a growth–defense tradeoff constrains habitat-associated phenotypic divergence

Negative phenotypic correlations between growth and defensive traits are widespread in plants (Herms and Mattson 1992; Cipollini 2004; Züst and Agrawal 2017). If these growth–defense tradeoffs are a consequence of resource allocation constraints, and selection on growth traits differs strongly between light habitats, tradeoffs may prevent local adaptation of herbivore resistance across the landscape. Observations from our common gardens suggest that any growth–defense tradeoffs that might exist in bittercress did not pose such a constraint. Plants from sun source sites were more resistant to *S. nigrita* than plants from shade source sites when both were grown in a common shade environment, even though plants from sun source sites concurrently expressed stronger SAS-related phenotypes under these conditions.

Our results are consistent with emerging evidence that a tradeoff between SAS and defense can be decoupled through mutation and natural selection (Züst and Agrawal 2017). Prolonged exposure to different regimes of herbivore pressure can alter correlations between traits involved in competitive growth and resistance to herbivory in experimentally evolved plant populations, despite evidence that resource allocation constraints ultimately limit the extent to which both suites of traits can be co-expressed (Uesugi et al. 2017). Functional genetic studies of *Arabidopsis* mutants provide plausible mechanisms for the uncoupling of growth and defense at the molecular level (Moreno et al. 2009; Robson et al. 2010; Keuskamp et al. 2010; Kazan and Manners 2011; Keller et al. 2011; Cerrudo et al. 2012). Intriguingly, differences in the plastic response of bittercress to light habitat in our common gardens mirror those observed among *Arabidopsis* lines with mutations introduced to genes involved in light-responsive signaling (see Supplementary Discussion). Future studies could test if habitat-associated divergence in bittercress growth and defense phenotypes has evolved through genetic or epigenetic mutations to the well-characterized genes that mediate growth–defense tradeoffs in response to light environment in *Arabidopsis*.

We note, however, that our results do not provide unambiguous evidence for a decoupling of a growth–defense tradeoff in bittercress. Tradeoffs that arise through pleiotropy or resource allocation constraints can be masked when genotypes vary in their efficiency of resource acquisition or use (Agrawal et al. 2010; Züst and Agrawal 2017). Although rhizome size was standardized in our common garden experiment, differences in the quality or quantity of stored resources in rhizomes collected from different habitat sites, combined with habitat-dependent effects of this resource variation on growth and resistance phenotypes, might explain why plants from sun sources exhibited strong SAS without attenuated resistance to *S. nigrita.*

### Potential mechanisms underlying microgeographic variation in bittercress

Future studies are needed to discern the mechanisms through which habitat-associated growth and resistance phenotypes are inherited in bittercress. Whether these traits are inherited through genetic or non-genetic mechanisms has implications for the ecological processes that generate the phenotype-habitat matching we observed across the landscape.

Transgenerational effects of parental environment on the phenotypes of clonally propagated offspring (Schwaegerle et al. 2000; Latzel and Klimešová 2010) and seed propagated offspring (Galloway 2005; Galloway and Etterson 2007) have been well documented in plants. These effects can shape offspring resistance against herbivory (Agrawal 2001; Steets and Ashman 2010; Rasmann et al. 2012; González et al. 2017; Colicchio 2017), and epigenetic variants induced by (and affecting) resistance phenotypes can persist for multiple generations (Rasmann et al. 2012). Habitat matching of heritable phenotypes in bittercress could therefore be driven simply by parental experience, without requiring natural selection to filter locally unfit genotypes from sun and shade habitats.

If heritable variation in the expression of growth and defense phenotypes has a genetic basis, however, matching of these phenotypes to local habitats would require strong, spatially varying selection to overcome homogenizing effects of gene flow across fine-grained habitat mosaics (Lenormand 2002). There is strong evidence that this filtering process can occur in plants, sometimes at the scale of only a few meters (Waser and Price 1989; Schmitt and Gamble 1990; Antonovics 2006; Schemske and Bierzychudek 2007; Hendrick et al. 2016).

By reducing the frequency of inter-habitat mating events, differences in flowering time among bittercress in sun and shade habitats, coupled with natural selection filtering unfit genotypes in each habitat, could facilitate the buildup of locally adaptive genetic variants or maternal effects (Levin 2009). We quantified flowering time between sun and shade habitats at two sites, and found a marked difference in one of these sites, although the duration of flowering time still overlapped among habitats. The observed phenological differences between sun and shade sites are one of many potential mechanisms that may reinforce divergence between sun and shade habitats in bittercress.

### Implications for the ecology and evolution of bittercress–*Scaptomyza* interactions

In a series of seminal field studies in the 1980s and 1990s, Louda and colleagues showed that higher herbivory on bittercress in open sun habitats, including by the abundant specialist herbivore *S. nigrita*, likely drives the distribution of bittercress toward more shaded habitats (Louda et al. 1987; Louda and Rodman 1996). These studies serve as a textbook example (Ricklefs and Miller 2000) of how natural enemies shape the distribution of their hosts across a heterogeneous landscape. However, the ultimate cause of higher herbivory in the sun remained unresolved.

Given that expression of SAS is frequently associated with reduced defenses against herbivory, and shade-derived bittercress exhibit reduced SAS relative to sun-derived bittercress in their source habitats, one possibility is that herbivory is higher in the sun because bittercress plants in the sun are more palatable. The fact that shade source plants are less resistant than sun source plants to *S. nigrita* larval herbivory does not support this interpretation. Instead, our results further support the conclusion, emerging from other recent work (Alexandre et al. 2017), that a preference of *S. nigrita* for sun-exposed habitats is likely sufficient to explain higher herbivory on bittercress in the sun. Thus, this textbook example of how natural enemies shape the distribution of their hosts is ultimately driven by the effect of an extrinsic habitat characteristic (light) on the behavior of an herbivore, not by the herbivore’s behavioral response to intrinsic differences in host plant quality across environments.

### Conclusion

We found that bittercress offspring, whether produced clonally or from seed, exhibit growth and defense phenotypes that match their parental habitat across a fine-grained (i.e., microgeographic) habitat mosaic. If this process leads offspring to have higher viability or fecundity in their natal habitats, it could be an important mechanism that reduces rates of inter-habitat recruitment and gene flow, in turn facilitating the maintenance of genetic and epigenetic variation within populations (Felsenstein 1976; Hedrick 2006). The ecological, physiological, and evolutionary mechanisms underlying phenotype-habitat matching in bittercress—and their generality across plant systems—await future study.

## ACKNOWLEDGMENTS

We acknowledge Ian Billick (RMBL), Jennifer Reithel (RMBL), Kailen Mooney (UC-Irvine), and Carol Boggs (University of South Carolina) for advice during the design and data collection phases of this project. Financial support was provided to N.K.W. by the RMBL (Research Fellowships 2010–2013), the National Science Foundation Division of Environmental Biology (NSF DEB-1256758), the John Templeton Foundation (41855) and the National Institute of General Medical Sciences of the National Institutes of Health under Award Number R35GM119816; to P.T.H. by RMBL Graduate Fellowships (2011–2013), the University of Arizona Center for Insect Science, and the NSF (DEB-1309493); to A.D.G. by the NSF (NSF DEB-1405966), an NSF Graduate Research Fellowship, and an RMBL Graduate Fellowship (2011); and to J.F. and S.F. by RMBL undergraduate research awards.

## ELECTRONIC SUPPLEMENTARY MATERIAL

**Fig. S1.**
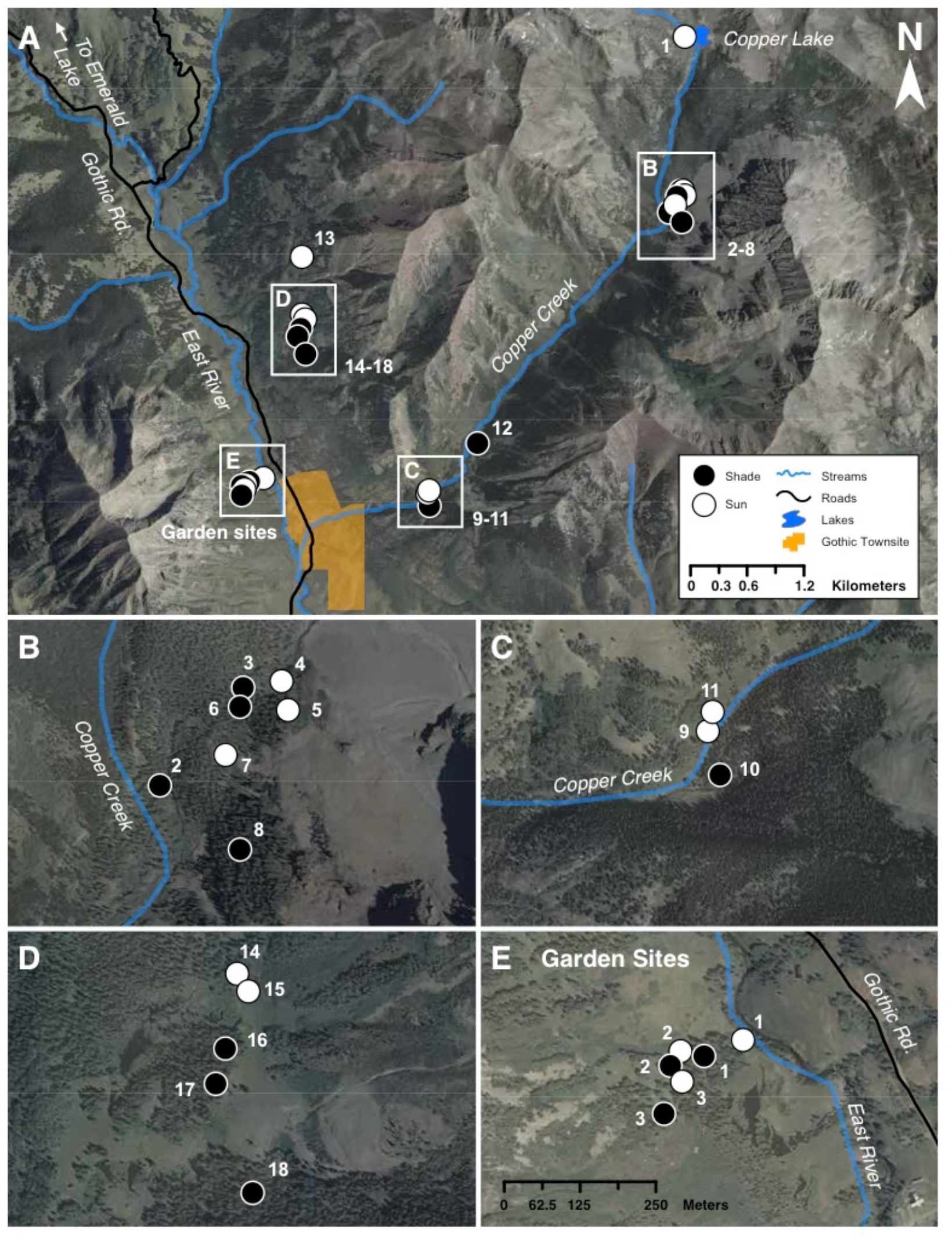
Map of source and garden sites used in the field common garden study in the East River Valley and Copper Creek drainages, near the RMBL in Gothic, CO. (**A)**. Base map showing all sites within region (1:48,000). (**B–E)**. Maps showing detail of site locations (all same scale, 1:7,500). Scale bar in E applies to B–E.

**Fig. S2.**
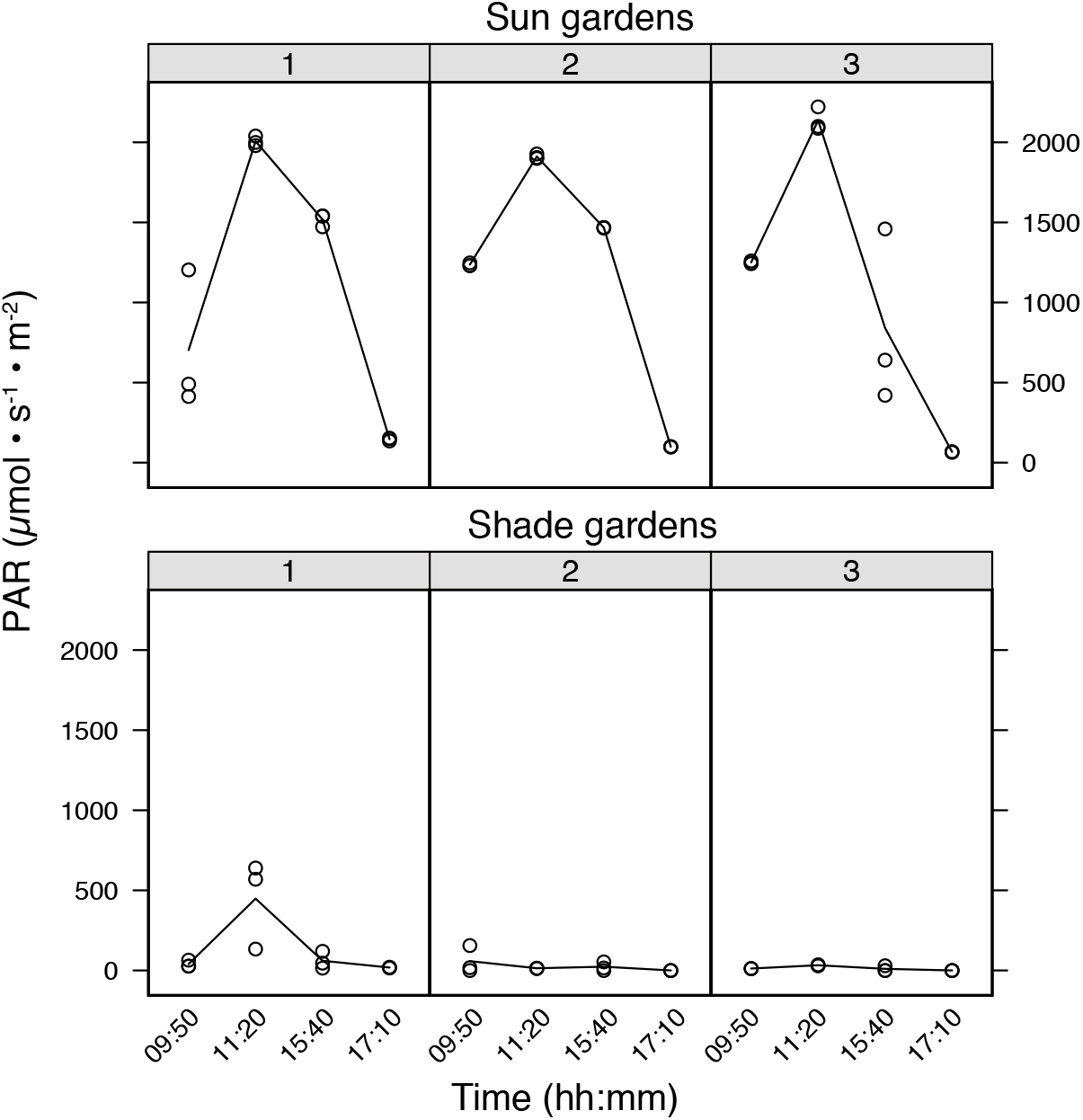
PAR measurements for the six common garden sites showing daily natural variation in light abundance. Line depicts mean of three measurements per time point per garden.

**Fig. S3.**
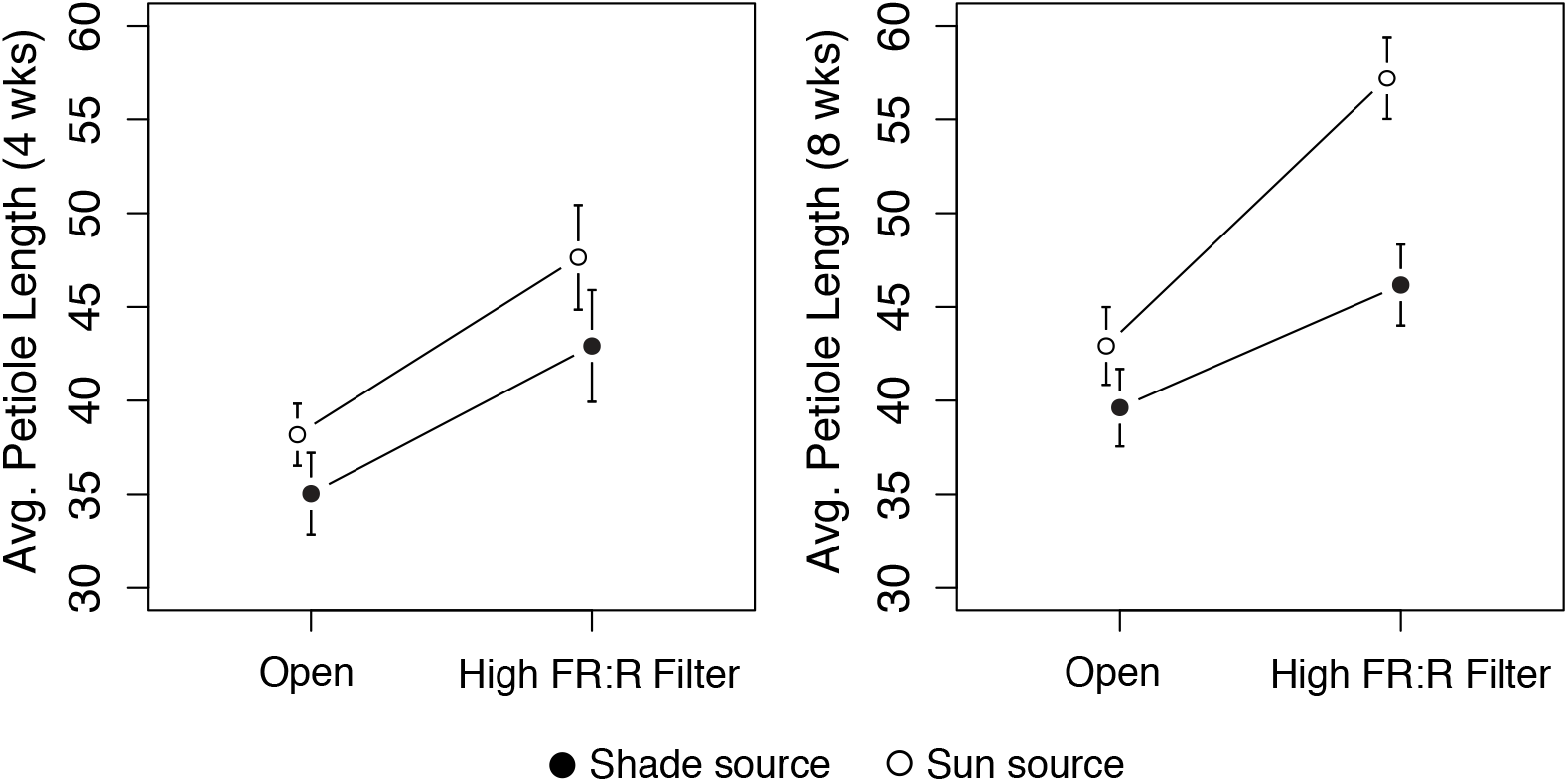
Petiole length of shade and sun source bittercress re-grown from seed under neutral (“Open”) or light filters (“High FR:R Filter”) to simulate neighbor shading in the greenhouse. ± indicates standard error.

**Fig. S4.**
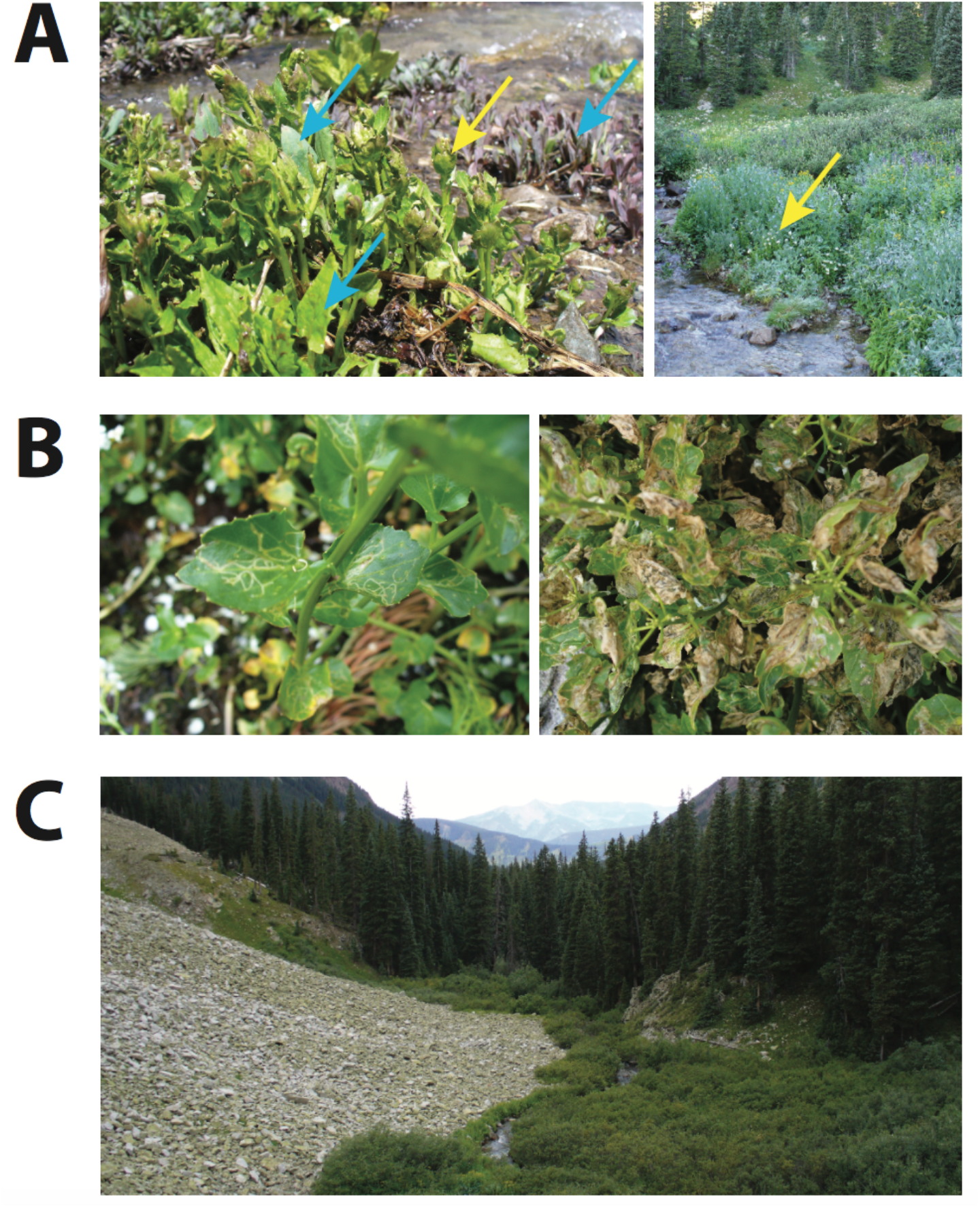
Photographs illustrating characteristics of open sun habitats for bittercress near RMBL. (**A**) Bittecress densely interspersed among heterospecific forbs early (left) and later (right) in the growing season. Yellow arrows: bittercress (with white flowers in the image to the right); blue arrows: other forbs. (**B**) Leaf mines from *S. nigrita* early (left) and later (right) in the growing season. (**C**) An alpine stream providing a patch of sunny habitat for bittercress at the base of a talus slope, before flowing into a shaded evergreen forest.

**Table S1.** Attributes of sites used as sources for bittercress genets for common gardens.

| Site # | Date Collected | Lat. | Long. | Elevation (m) | Soil Moisture | Light Env. | PAR ($\mu\text{mol} \cdot \text{s} \cdot \text{m}^{-2}$)<sup>1</sup> | % Canopy Open | Average DBH (cm) |
| --- | --- | --- | --- | --- | --- | --- | --- | --- | --- |
| 1 | 10-Jul-11 | 39.0050602776535 | -106.943317166893 | 3461 | Very Wet | Sun | 1980 | 99.84 | 0 |
| 2 | 10-Jul-11 | 38.9885304491745 | -106.945774321867 | 3218 | Moist Loamy | Shade | 300 | 6.76 | 10 |
| 3 | 11-Jul-11 | 38.9900162385746 | -106.944210201638 | 3240 | Moist Loamy | Shade | 130 | 8.06 | 121.75 |
| 4 | 11-Jul-11 | 38.9900829890862 | -106.943450029132 | 3253 | Very Wet | Sun | 2030 | 82.94 | 0 |
| 5 | 11-Jul-11 | 38.9896982273227 | -106.9432894133 | 3261 | Very Wet | Sun | 2034 | 95.42 | 0 |
| 6 | 11-Jul-11 | 38.9897805369006 | -106.94429611421 | 3244 | Moist Loamy | Shade | 130 | 4.16 | 83.25 |
| 7 | 11-Jul-11 | 38.9890385630266 | -106.944483622637 | 3220 | Very Wet | Sun | 2134 | 99.84 | 0 |
| 8 | 11-Jul-11 | 38.9876108211294 | -106.944190554921 | 3250 | Moist Loamy | Shade | 28 | 3.38 | 81.75 |
| 9 | 12-Jul-11 | 38.9607056556777 | -106.973679741745 | 3023 | Very Wet | Sun | 2026 | 88.14 | 0 |
| 10 | 12-Jul-11 | 38.9600423351315 | -106.973476684125 | 3013 | Wet Loamy | Shade | 35 | 5.72 | 72.25 |
| 11 | 12-Jul-11 | 38.9609507737634 | -106.973571138078 | 3994 | Very Wet | Sun | 1933 | 86.84 | 0 |
| 12 | 12-Jul-11 | 38.9655594715881 | -106.968516958823 | 3994 | Very Wet | Shade | 32 | 3.9 | 101 |
| 13 | 13-Jul-11 | 38.9828665210329 | -106.989822099201 | 4058 | Very Wet | Sun | 1780 | 96.2 | 0 |
| 14 | 13-Jul-11 | 38.9774145794955 | -106.98986568487 | 4082 | Very Wet | Sun | 1870 | 88.4 | 0 |
| 15 | 13-Jul-11 | 38.9771831850899 | -106.989697600334 | 4057 | Very Wet | Sun | 1850 | 99.84 | 0 |
| 16 | 13-Jul-11 | 38.976356744414 | -106.990078474138 | 4052 | Very Wet | Shade | 54 | 4.68 | 98 |
| 17 | 13-Jul-11 | 38.975757817399 | -106.990327194402 | 4046 | Moist Loamy | Shade | 60 | 4.16 | 140.5 |
| 18 | 13-Jul-11 | 38.974195183581 | -106.989498527449 | 4051 | Very Wet | Shade | 40 | 7.28 | 78.75 |

<sup>1</sup>Light meter was positioned above the center of each source collection plot, at times without cloud cover between 1:00 pm and 3:00 pm during the week of July 17, 2011.

**Table S2.** Environmental attributes of sun and shade source sites used in field common gardens.

| Attribute | Sun Sites (N = 9) |  | Shade sites (N = 9) |  | ANOVA |  |
| --- | --- | --- | --- | --- | --- | --- |
| | $\mu$ | se | $\mu$ | se | *F* | *P* |
| PAR ($\mu\text{mol}\cdot\text{s}^{-1}\cdot\text{m}^{-2}$) | 1959.7 | 37.0 | 89.0 | 29.4 | 1564 | <10<sup>-10</sup> |
| % Canopy Open | 93.1 | 2.2 | 5.3 | 0.6 | 1527 | <10<sup>-10</sup> |
| DBH (cm) | 0.00 | 0.00 | 87.5 | 12.2 | 51.77 | <10<sup>-10</sup> |
| Elevation (m) | 3601 | 146 | 3568 | 150 | 0.025 | 0.875 |

**Table S3.**
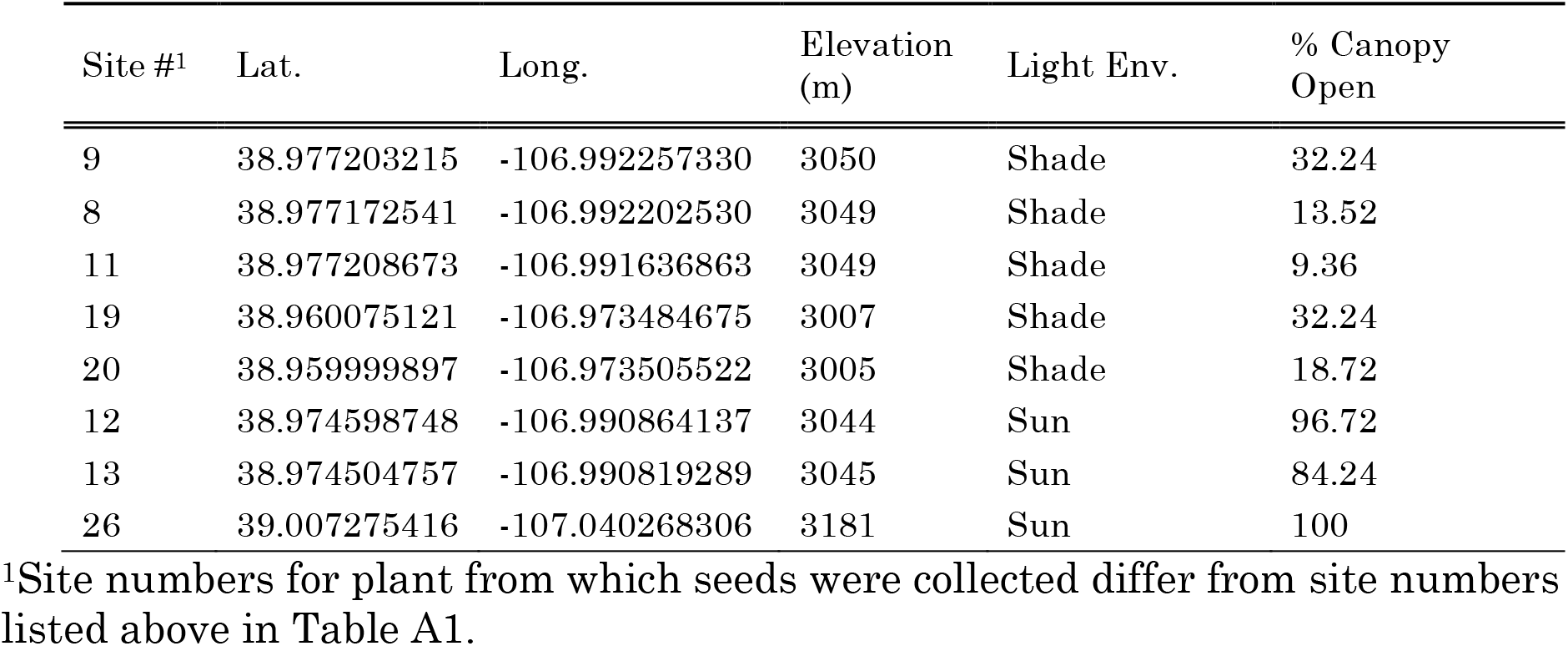
Source site attributes for plants from which seeds were collected and regrown for use in greenhouse common garden experiment.

**Table S4.**
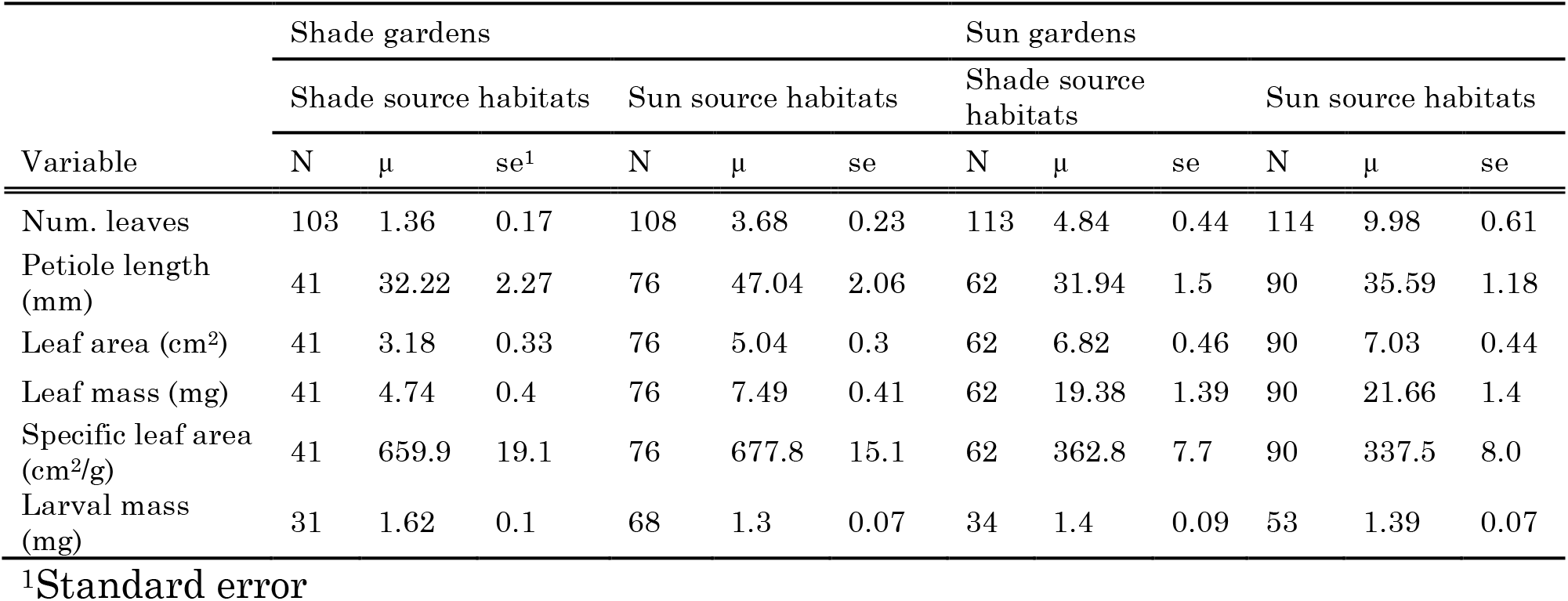
Descriptive statistics for plant growth and herbivory phenotypes measured in common gardens.

| Variable | Shade gardens |  |  |  |  |  | Sun gardens |  |  |  |  |  |
| --- | --- | --- | --- | --- | --- | --- | --- | --- | --- | --- | --- | --- |
|  | Shade source habitats |  |  | Sun source habitats |  |  | Shade source habitats |  |  | Sun source habitats |  |  |
|  | N | μ | se<sup>1</sup> | N | μ | se | N | μ | se | N | μ | se |
| Num. leaves | 103 | 1.36 | 0.17 | 108 | 3.68 | 0.23 | 113 | 4.84 | 0.44 | 114 | 9.98 | 0.61 |
| Petiole length (mm) | 41 | 32.22 | 2.27 | 76 | 47.04 | 2.06 | 62 | 31.94 | 1.5 | 90 | 35.59 | 1.18 |
| Leaf area (cm<sup>2</sup>) | 41 | 3.18 | 0.33 | 76 | 5.04 | 0.3 | 62 | 6.82 | 0.46 | 90 | 7.03 | 0.44 |
| Leaf mass (mg) | 41 | 4.74 | 0.4 | 76 | 7.49 | 0.41 | 62 | 19.38 | 1.39 | 90 | 21.66 | 1.4 |
| Specific leaf area (cm<sup>2</sup>/g) | 41 | 659.9 | 19.1 | 76 | 677.8 | 15.1 | 62 | 362.8 | 7.7 | 90 | 337.5 | 8.0 |
| Larval mass (mg) | 31 | 1.62 | 0.1 | 68 | 1.3 | 0.07 | 34 | 1.4 | 0.09 | 53 | 1.39 | 0.07 |

<sup>1</sup>Standard error

### Supplementary Discussion

Functional genetic studies of *Arabidopsis* mutants provide plausible, although speculative, mechanisms for the uncoupling of growth and defense at the molecular level in bittercress. Recall that bittercress from shade source habitats expressed relatively reduced SAS as well as reduced herbivore resistance when grown in shade gardens. This pattern was found in the SAS-deficient *A. thaliana* mutant *sav3-2,* which exhibits an attenuated defense phenotype but no SAS expression under high FR:R light (Moreno et al. 2009; Cerrudo et al. 2012). It is possible that bittercress in evergreen forest canopy understories do not *express* SAS but continue to express attenuated resistance to herbivores through a pleiotropic effect associated with detection of high FR:R light. In sun gardens, shade-derived bittercress resisted herbivores to the same degree as plants from sun source. Decoupling of shade detection from SAS expression would allow plants adapted to evergreen forest canopy shade to continue to resist herbivores in sun habitats while reducing the expression of SAS following shade detection under evergreen forest canopy shade, where its expression is unnecessary.

In contrast, plants from sun habitats could activate SAS upon detection of high FR:R light but may have decoupled the resistance attenuation that typically results from detection of neighbor shade (Robson et al. 2010; Kazan and Manners 2011). This could occur via modification of the physiological basis of shade detection. In *A. thaliana*, SAS is also induced by depletion of blue (B) light, which is detected by cryptochrome 1 (CRY1) (Keuskamp et al. 2010; Keller et al. 2011). Mutant *A. thaliana* with inactivated CRY1 constitutively express features of SAS but without attenuation of JA-dependent defenses. The same phenotype is produced in wild-type *A. thaliana* grown under B-depleted light (Cerrudo et al. 2012), suggesting that alternate light-sensing pathways may interact to determine the consequences of shade detection for plant defense against herbivores.

Mutations that reduce the ecological costs of negative correlations may be favored under particular combinations of selective agents, such as those faced by bittercress in sun habitats. Novel and habitat-specific genotypes may emerge that reveal novel insights into the genetic architecture of phenotypic plasticity. Extensive quantitative genetic variation has been found for SAS expression among *A. thaliana* accessions (Filiault and Maloof 2012), and testing for correlations between SAS and defense-related genetic variation coupled to shading and herbivory experiments may be one promising avenue of future research.

## References

Abràmoff MD, Magalhães PJ, Ram SJ (2004). Image processing with ImageJ. Biophoton Int 11:36–42 doi:10.1002/9780470089941.eta03cs9.

Agrawal AA, Hastings AP, Johnson MT, Maron JL, Salminen J (2012). Insect herbivores drive real-time ecological and evolutionary change in plant populations. Science 338:113–116 doi: 10.1126/science.1225977

Agrawal AA, Conner JK, Rasmann S (2010). Tradeoffs and negative correlations in evolutionary ecology. In: M.A. Bell, W.F. Eanes, D.J. Futuyma, and J.S. Levinton, eds. Evolution After Darwin: the First 150 Years. Sinauer Associates, Sunderland, MA.

Agrawal AA (2001). Transgenerational consequences of plant responses to herbivory: An adaptive maternal effect? Am Nat 157:555–569 doi: 10.1086/319932

Alexandre NM, Humphrey PT, Frazier J, Gloss AD, Lee J, Affeldt HA, Whiteman NK (2017). Habitat preference of an herbivore shapes the habitat distribution of its host plant. bioRxiv:156240 doi: 10.1101/156240

Alsdurf JD, Ripley TJ, Matzner SL, Siemens DH (2013). Drought-induced trans-generational tradeoff between stress tolerance and defence: Consequences for range limits? AoB Plants 5: plt038 doi: 10.1093/aobpla/plt038

Antonovics J (2006). Evolution in closely adjacent plant populations X: Long-term persistence of prereproductive isolation at a mine boundary. Heredity 97:33 doi: 10.1038/sj.hdy.6800835

Bates D, Maechler M, Bolker B, Walker S (2014). lme4: Linear mixed-effects models using Eigen and S4. R package v1.7: 1–23

Bell DL, Galloway LF (2008). Population differentiation for plasticity to light in an annual herb: Adaptation and cost. Am J Bot 95:59–65 doi: 10.3732/ajb.95.1.59

Boege K (2010). Induced responses to competition and herbivory: Natural selection on multi-trait phenotypic plasticity. Ecology 91:2628–2637

Brachi B, Meyer CG, Villoutreix R, Platt A, Morton TC, Roux F, Bergelson J (2015). Coselected genes determine adaptive variation in herbivore resistance throughout the native range of arabidopsis thaliana. Proc Natl Acad Sci U S A 112:4032–4037 doi: 10.1073/pnas.1421416112

Cerrudo I, Keller MM, Cargnel MD, Demkura PV, de Wit M, Patitucci MS, Pierik R, Pieterse CM, Ballare CL (2012). Low red/far-red ratios reduce Arabidopsis resistance to Botrytis cinerea and jasmonate responses via a COI1-JAZ10-dependent, salicylic acid-independent mechanism. Plant Physiol 158:2042–2052 doi: 10.1104/pp.112.193359

Cipollini D (2004). Stretching the limits of plasticity: Can a plant defend against both competitors and herbivores? Ecology 85:28–37 doi: 10.1890/02-0615

Colicchio J (2017). Transgenerational effects alter plant defence and resistance in nature. J Evol Biol 30:664–680 doi: 10.1111/jeb.13042

Collinge SK, Louda SM (1989). Scaptomyza nigrita Wheeler (diptera: Drosophilidae), a leaf miner of the native crucifer, Cardamine cordifolia A. gray (bittercress). J. Kans. Entomol. Soc. 62:1–10

Donohue K, Hammond Pyle E, Messiqua D, Shane Heschel M, Schmitt J (2001). Adaptive divergence in plasticity in natural populations of Impatiens capensis and its consequences for performance in novel habitats. Evolution 55:692–702 doi: 10.1554/0014-3820(2001)055[0692:ADIPIN]2.0.CO;2

Donohue K, Messiqua D, Pyle EH, Heschel MS, Schmitt J (2000). Evidence of adaptive divergence in plasticity: Density- and site-dependent selection on shade-avoidance responses in Impatiens capensis. Evolution 54:1956–1968 doi: 10.1111/j.0014-3820.2000.tb01240.x

Dostálek T, Rokaya MB, Maršík P, Rezek J, Skuhrovec J, Pavela R, Münzbergová Z (2016). Trade-off among different anti-herbivore defence strategies along an altitudinal gradient. AoB Plants 8: plw026 doi: 10.1093/aobpla/plw026

Dudley S, Schmitt J (1995). Genetic differentiation in morphological responses to simulated foliage shade between populations of Impatiens capensis from open and woodland sites. Funct Ecol 9:655–666 doi: 10.2307/2390158

Dudley SA, Schmitt J (1996). Testing the adaptive plasticity hypothesis: Density-dependent selection on manipulated stem length in Impatiens capensis. Am Nat 147:445–465 doi: 10.1086/285860

Felsenstein J (1976). The theoretical population genetics of variable selection and migration. Annu Rev Genet 10:253–280 doi: 10.1146/annurev.ge.10.120176.001345

Galen C, Shore JS, Deyoe H (1991). Ecotypic divergence in alpine *Polemonium viscosum*: Genetic structure, quantitative variation, and local adaptation. Evolution 45:1218–1228

Galloway LF (2005). Maternal effects provide phenotypic adaptation to local environmental conditions. New Phytol 166:93–100 doi: 10.1111/j.1469-8137.2004.01314.x

Galloway LF, Etterson JR (2007). Transgenerational plasticity is adaptive in the wild. Science 318:1134–1136 doi: 10.1126/science.1148766

Gloss AD, Vassao DG, Hailey AL, Nelson Dittrich AC, Schramm K, Reichelt M, Rast TJ, Weichsel A, Cravens MG, Gershenzon J, Montfort WR, Whiteman NK (2014). Evolution in an ancient detoxification pathway is coupled with a transition to herbivory in the Drosophilidae. Mol Biol Evol 31:2441–2456 doi: 10.1093/molbev/msu201

González APR, Dumalasová V, Rosenthal J, Skuhrovec J, Latzel V (2017). The role of transgenerational effects in adaptation of clonal offspring of white clover (*Trifolium repens*) to drought and herbivory. Evol Ecol 31:345–361 doi: 10.1007/s10682-016-9844-5

Greig-Smith P (1952). The use of random and contiguous quadrats in the study of the structure of plant communities. Annals of Botany 16:293–316 doi: 10.1093/oxfordjournals.aob.a083317

Haldane JBS (1930). A mathematical theory of natural and artificial selection (part VI, isolation). Proceedings of the Cambridge Philosophical Society 26:220–230 doi: 10.1017/S0305004100015450

Halekoh U, Højsgaard S (2014). A Kenward-Roger approximation and parametric bootstrap methods for tests in linear mixed models–the R package pbkrtest. Journal of Statistical Software 59: 1–32 doi: 10.18637/jss.v059.i09.

Hedrick PW (2006). Genetic polymorphism in heterogeneous environments: The age of genomics. Annu Rev Ecol Evol Syst 37:67–93 DOI 10.1146/annurev.ecolsys.37.091305.110132

Hendrick MF, Finseth FR, Mathiasson ME, Palmer KA, Broder EM, Breigenzer P, Fishman L (2016). The genetics of extreme microgeographic adaptation: An integrated approach identifies a major gene underlying leaf trichome divergence in Yellowstone Mimulus guttatus. Mol Ecol 25:5647–5662 doi: 10.1111/mec.13753

Herms DA, Mattson WJ (1992). The dilemma of plants: To grow or defend. Q Rev Biol 67:283–335 doi: 10.1086/417659

Humphrey PT, Gloss AD, Alexandre NM, Villalobos MM, Fremgen MR, Groen SC, Meihls LN, Jander G, Whiteman NK (2016). Aversion and attraction to harmful plant secondary compounds jointly shape the foraging ecology of a specialist herbivore. Ecology and evolution 6:3256–3268 doi: 10.1002/ece3.2082

Humphrey PT, Nguyen TT, Villalobos MM, Whiteman NK (2014). Diversity and abundance of phyllosphere bacteria are linked to insect herbivory. Mol Ecol 23:1497–1515 doi: 10.1111/mec.12657

Kazan K, Manners JM (2011). The interplay between light and jasmonate signalling during defence and development. J Exp Bot 62:4087–4100 doi: 10.1093/jxb/err142

Kenward MG, Roger JH (1997). Small sample inference for fixed effects from restricted maximum likelihood. Biometrics 53:983–997 doi: 0.2307/2533558.

Keller MM, Jaillais Y, Pedmale UV, Moreno JE, Chory J, Ballaré CL (2011). Cryptochrome 1 and phytochrome B control shade-avoidance responses in *Arabidopsis* via partially independent hormonal cascades. The Plant Journal 67:195–207 doi: 10.1111/j.1365-313X.2011.04598.x

Keuskamp DH, Sasidharan R, Pierik R (2010). Physiological regulation and functional significance of shade avoidance responses to neighbors. Plant Signaling & Behavior 5:655–662 doi: 10.4161/psb.5.6.11401

Latzel V, Klimešová J (2010). Transgenerational plasticity in clonal plants. Evolutionary Ecology 24: 1537–1543 doi: 10.1007/s10682-010-9385-2

Lenormand T (2002). Gene flow and the limits to natural selection. Trends in Ecology & Evolution 17:183–189 doi: 10.1016/S0169-5347(02)02497-7

Levin DA (2009). Flowering-time plasticity facilitates niche shifts in adjacent populations. New Phytol 183:661–666 doi: 10.1111/j.1469-8137.2009.02889.x

Linhart YB, Grant MC (1996). Evolutionary significance of local genetic differentiation in plants. Annu Rev Ecol Syst 27:237–277 doi: 10.1146/annurev.ecolsys.27.1.237

Louda SM, Rodman JE (1996). Insect herbivory as a major factor in the shade distribution of a native crucifer (*Cardamine cordifolia* A. Gray, bittercress). J Ecol 84:229–237 doi: 10.2307/2261358

Louda SM (1984). Herbivore effect on stature, fruiting, and leaf dynamics of a native crucifer. Ecology 65:1379–1386 doi: 10.2307/1939118

Louda SM, Dixon PM, Huntly NJ (1987). Herbivory in sun versus shade at a natural meadow-woodland ecotone in the Rocky Mountains. Plant Ecol 72:141–149

Moreno JE, Tao Y, Chory J, Ballare CL (2009). Ecological modulation of plant defense via phytochrome control of jasmonate sensitivity. Proc Natl Acad Sci U S A 106:4935–4940 doi: 10.1073/pnas.0900701106

Pellissier L, Roger A, Bilat J, Rasmann S (2014). High elevation *Plantago lanceolata* plants are less resistant to herbivory than their low elevation conspecifics: Is it just temperature? Ecography 37:950–959 doi: 10.1111/ecog.00833

Prasad KV, Song BH, Olson-Manning C, Anderson JT, Lee CR, Schranz ME, Windsor AJ, Clauss MJ, Manzaneda AJ, Naqvi I, Reichelt M, Gershenzon J, Rupasinghe SG, Schuler MA, Mitchell-Olds T (2012). A gain-of-function polymorphism controlling complex traits and fitness in nature. Science 337:1081–1084 10.1126/science.1221636

R Core Team (2013). R: A language and environment for statistical computing. R Foundation for Statistical Computing, Vienna, Austria. v3.3. URL http://www.R-project.org/

Rasmann S, De Vos M, Casteel CL, Tian D, Halitschke R, Sun JY, Agrawal AA, Felton GW, Jander G (2012). Herbivory in the previous generation primes plants for enhanced insect resistance. Plant Physiol 158:854–863 doi 10.1104/pp.111.187831

Richardson JL, Urban MC, Bolnick DI, Skelly DK (2014). Microgeographic adaptation and the spatial scale of evolution. Trends in Ecology & Evolution 29:165–176 doi: 10.1016/j.tree.2014.01.002

Ricklefs R, Miller G (2000). Ecology, 4th edn. W.H. Freeman, New York

Robson F, Okamoto H, Patrick E, Harris SR, Wasternack C, Brearley C, Turner JG (2010). Jasmonate and phytochrome A signaling in *Arabidopsis* wound and shade responses are integrated through JAZ1 stability. Plant Cell 22:1143–1160 doi 10.1105/tpc.109.067728

Runkle ES, Heins RD (2001). Specific functions of red, far red, and blue light in flowering and stem extension of long-day plants. J Am Soc Hort Sci 126:275–282

Sato Y, Kudoh H (2017). Fine-scale frequency differentiation along a herbivory gradient in the trichome dimorphism of a wild Arabidopsis. Ecology and evolution 7:2133–2141 doi: 10.1002/ece3.2830

Schemske DW, Bierzychudek P (2007). Spatial differentiation for flower color in the desert annual linanthus parryae: Was wright right? Evolution 61:2528–2543 doi: 10.1111/j.1558-5646.2007.00219.x

Schmitt J, McCormac AC, Smith H (1995). A test of the adaptive plasticity hypothesis using transgenic and mutant plants disabled in phytochrome-mediated elongation responses to neighbors. Am Nat 146:937–953 doi: 10.1086/285832

Schmitt J, Gamble SE (1990). The effect of distance from the parental site on offspring performance and inbreeding depression in *Impatiens capensis*: A test of the local adaptation hypothesis. Evolution 44:2022–2030 doi: 10.1111/j.1558-5646.1990.tb04308.x

Schwaegerle KE, McIntyre H, Swingley C (2000). Quantitative genetics and the persistence of environmental effects in clonally propagated organisms. Evolution 54:452–461 doi: 10.1111/j.0014-3820.2000.tb00048.x

Siemens DH, Haugen R (2013). Plant chemical defense allocation constrains evolution of tolerance to community change across a range boundary. Ecology and Evolution 3:4339–4347 doi: 10.1002/ece3.657

Siemens DH, Haugen R, Matzner S, Vanasma N (2009). Plant chemical defence allocation constrains evolution of local range. Mol Ecol 18:4974–4983 doi: 10.1111/j.1365-294X.2009.04389.x

Sork VL, Stowe KA, Hochwender C (1993). Evidence for local adaptation in closely adjacent subpopulations of northern red oak (*Quercus rubra* L.) expressed as resistance to leaf herbivores. Am Nat 142:928–936 doi: 10.1086/285581

Steets JA, Ashman T (2010). Maternal effects of herbivory in Impatiens capensis. Int J Plant Sci 171:509–518 doi: 10.1086/651944

Uesugi A, Connallon T, Kessler A, Monro K (2017). Relaxation of herbivore-mediated selection drives the evolution of genetic covariances between plant competitive and defense traits. Evolution 71:1700–1709 doi: 10.1111/evo.13247

Uriarte M, Canham CD, Root RB (2002). A model of simultaneous evolution of competitive ability and herbivore resistance in a perennial plant. Ecology 83:2649–2663 doi: 10.2307/3072004

Valladares F, Gianoli E, Gómez JM (2007). Ecological limits to plant phenotypic plasticity. New Phytol 176:749–763 doi: 10.1111/j.1469-8137.2007.02275.x

Van Kleunen M, Fischer M (2005). Constraints on the evolution of adaptive phenotypic plasticity in plants. New Phytol 166:49–60 doi: 10.1111/j.1469-8137.2004.01296.x

Venables WN, Ripley BD (2002). Modern Applied Statistics with S. 4th edn. Springer, New York

Waser NM, Price MV (1989). Optimal outcrossing in *Ipomopsis aggregata*: seed set and offspring fitness. Evolution 43:1097–1109 doi: 10.2307/2409589

Whiteman NK, Gloss AD, Sackton TB, Groen SC, Humphrey PT, Lapoint RT, Sonderby IE, Halkier BA, Kocks C, Ausubel FM, Pierce NE (2012). Genes involved in the evolution of herbivory by a leaf-mining, drosophilid fly. Genome Biol Evol 4:900–916 doi: 10.1093/gbe/evs063

Whiteman NK, Groen SC, Chevasco D, Bear A, Beckwith N, Gregory TR, Denoux C, Mammarella N, Ausubel FM, Pierce NE (2011). Mining the plant-herbivore interface with a leafmining *Drosophila* of *Arabidopsis*. Mol Ecol 20:995–1014 doi: 10.1111/j.1365-294X.2010.04901.x

Züst T, Agrawal AA (2017). Trade-offs between plant growth and defense against insect herbivory: An emerging mechanistic synthesis. Annual Review of Plant Biology 68:513–534 doi: 10.1146/annurev-arplant-042916-040856

Züst T, Heichinger C, Grossniklaus U, Harrington R, Kliebenstein DJ, Turnbull LA (2012). Natural enemies drive geographic variation in plant defenses. Science 338:116–119 doi 10.1126/science.1226397

## References

Cerrudo I., M.M. Keller, M.D. Cargnel, P.V. Demkura, M. de Wit, M.S. Patitucci, R. Pierik, C.M. Pieterse, C.L. Ballare. (2012). Low red/far-red ratios reduce Arabidopsis resistance to Botrytis cinerea and jasmonate responses via a COI1-JAZ10-dependent, salicylic acid-independent mechanism. Plant Physiol. 158:2042–2052.

Filiault D. L., and J. N. Maloof. 2012. A genome-wide association study identifies variants underlying the Arabidopsis thaliana shade avoidance response. PLoS Genetics 8:e1002589.

Kazan K., J.M. Manners. (2011). The interplay between light and jasmonate signalling during defence and development. J. Exp. Bot. 62:4087–4100.

Keller M.M., Y. Jaillais, U.V. Pedmale, J.E. Moreno, J. Chory, C.L. Ballaré. (2011). Cryptochrome 1 and phytochrome B control shade-avoidance responses in *Arabidopsis* via partially independent hormonal cascades. The Plant Journal 67:195–207.

Keuskamp D. H., R. Sasidharan, and R. Pierik. 2010. Physiological regulation and functional significance of shade avoidance responses to neighbors. Plant signaling & behavior 5:655–662.

Moreno J. E., Y. Tao, J. Chory, and C. L. Ballare. 2009. Ecological modulation of plant defense via phytochrome control of jasmonate sensitivity. Proc. Natl. Acad. Sci. U. S. A. 106:4935–4940.

Robson F., H. Okamoto, E. Patrick, S. R. Harris, C. Wasternack, C. Brearley, and J. G. Turner. 2010. Jasmonate and phytochrome A signaling in Arabidopsis wound and shade responses are integrated through JAZ1 stability. Plant Cell 22:1143–1160.

